# Evaluation of rearing parameters of a self-limiting strain of the Mediterranean fruit fly, *Ceratitis capitata* (Diptera: Tephritidae)

**DOI:** 10.1101/404749

**Authors:** Elaini Rachid, Romisa Asadi, Neil Naish, Martha Koukidou, Mazih Ahmed

## Abstract

The Mediterranean fruit fly, *Ceratitis capitata*, (medfly) is an important pest of stone and pome fruit, causing significant economic losses worldwide. Current control is primarily based on insecticides, often mixed with protein baits. Chemical approaches are effective but there are label limits to avoid residues in fruits and harm to the environment and sustained use will lead to pesticide resistance in the medfly pest. In recent years, emphasis has been placed on environmentally friendly methods to control medfly.

Oxitec has developed a self-limiting medfly strain (OX3864A) that demonstrates conditional female-specific mortality in the early life stages. Sustained release of OX3864A males offers a mating-based approach to medfly control, which should lead to significant economic benefits in area-wide programmes. Furthermore, a heritable fluorescent marker provides quick and accurate identification of released OX3864A males for efficient monitoring in the field.

An important prerequisite of mating-based control programmes is the availability of adequate numbers of high-quality male flies in a sustainable and cost-effective manner. This paper summarises rearing optimisations for the OX3864A strain and the production of OX3864A males.

## Introduction

The Mediterranean fruit fly, *Ceratitis capitata* Wiedemann (Diptera: Tephritidae) (hereafter medfly), is an important pest of stone and pome fruit worldwide, particularly in tropical, sub-tropical and temperate climates (Gasperi et al. 2002), due to its great capacity for dispersion, adaptability and high rate of reproduction (Malacrida et al. 2007, Diamantidis et al. 2011). In addition to the significant losses in fruit yield and quality (Diamantidis et al. 2008), medfly is responsible for the establishment of expensive quarantine measures that prevent or hinder agricultural exports from medfly-endemic countries (Rendon et al. 2006).

Insecticide sprays have been the principal tool for medfly control globally. Public concern regarding health and environmental impacts of insecticide use, loss through de-registration of some insecticides and the development of insecticide resistance (Kakani and Mathiopoulos 2008, Grzywacz et al. 2010, Sluder et al. 2012, Kakani et al. 2014), have driven the development of sustainable approaches towards controlling this insect pest. Mass-trapping (Leza et al. 2008), mating disruption technologies (Welter et al. 2005), and the Sterile Insect Technique (SIT) (Hendrichs et al. 2002) have also been employed for the management of medfly. Of these, SIT has enabled local eradication, prevention and suppression of the medfly (Hendrichs 2000, Barry et al. 2002, Hendrichs et al. 2002, Suckling et al. 2014). Examples of eradication successes of other species using SIT include the New World screwworm, *Cochliomyia hominivorax*, from North and Central America and from Libya (Concha et al. 2016); the tsetse fly, *Glossina austeni*, from Unguja Island in Zanzibar, Tanzania (FAO/IAEA 2008); and the melon fly, *Bactrocera cucurbitae*, from Japan (Koyama et al. 2004).

The SIT relies on the mass-production of factory-reared insects and their subsequent sterilisation through irradiation. Sterile insects are released *en masse* and on a sustained basis in the environment to achieve appropriate overflooding ratios. The sterile males mate with local wild females, leading to no or few viable offspring and the eventual reduction of the pest population. Higher overflooding ratios may result in local eradication (Hendrichs et al. 1995). SIT implementation requires the development of a cost-effective mass-rearing system in place (Vera et al. 2007), including cost-efficient larval and adult diets (Chan et al. 1990, Rehman et al. 2009, Ben-Yosef et al. 2015).

Irradiation is known to induce dominant lethal genetic effects in the male germline, which render males unable to produce viable progeny, but irradiation also provoke damage to somatic cells, resulting in reduced male competitiveness and higher operational costs (Barry et al. 2002, 2003, Lux et al. 2002, Kraaijeveld and Chapman 2004, San Andrés et al. 2007, Guerfali et al. 2011, Lauzon and Potter 2012, Rull et al. 2012).

The release of insects with ‘self-limiting’ genetic traits that induce early life-stage mortality to progeny, has been demonstrated as an effective management tool for specific insects across both agricultural and public health sectors (Harris et al. 2011, Ant et al. 2012, Lacroix et al. 2012, Harvey-Samuel et al. 2015, Gorman et al. 2016). The female-specific self-limiting approach allows for a male-only release cohort, that has been previously shown to increase the effectiveness of mating-based insect pest management programmes (Rendon et al. 2004). Mating between self-limiting males and wild females produces no viable female offspring, thereby decreasing the wild population. The addition of an antidote to the larval rearing medium prevents the expression of the self-limiting gene, allowing for the normal propagation of the strain.

The medfly strain OX3864A, genetically engineered to carry a conditional female-specific self-limiting gene, demonstrates full penetrance and benefits from an inherited genetic marker that enables quick and accurate identification from its wild counterparts. The potential of the sustained release of OX3864A males to control wild-type medfly populations has been previously demonstrated (Leftwich et al. 2014).

A vital prerequisite to any mating-based insect control programme is the optimal rearing of the insect strain for propagation and more importantly to produce the male-only release cohort. In this work, we evaluate the optimum quantity of mature and immature stages of the OX3864A strain and provide rearing and quality control parameters associated with the strain.

## Materials and Methods

### Adult colony maintenance

The OX3864A strain was maintained at the insect-rearing facilities of the Omnium Agricole du Souss Auxiliary Production Site, Chtouka, Morocco. All medfly life stages were maintained in a controlled environment (Relative Humidity-RH: 60-65%, 12:12 light:dark cycle), with adults reared at 25±1°C and larvae at 28±1°C.

Adults were reared in 77 cm × 72 cm × 7 cm wooden cages with 1:4 mixture of enzymatic yeast hydrolysate (Biokar diagnostics, A1202HA) and sugar as a rearing medium, and water containing tetracycline (Sigma, T3383) at a concentration of 100 mg/L. The cages were suspended on metallic frames, each of them holding approximately 11 cages. Cages were stocked with a known volume of pupae of the same age.

### Egg and pupal production

Eggs were oviposited through a fine mesh on the side wall of each cage and were collected in troughs of water beneath the cages, during a 24-h period. The egg-water solution was passed through a fine sieve and the retained eggs were washed off with a water bottle to a volumetric cylinder. Following volume measurement, eggs were transferred onto a damp 125 mm filter paper inside a 90-mm Petri dish until developed embryos could be identified.

Each egg cohort was seeded onto a kilogram of larval rearing medium as described by (Tanaka et al. 1969), with added tetracycline at a concentration of 100 ml/L for strain maintenance (hereafter Tet+), or without additives for male-only cohorts (male-only releases) (hereafter Tet-).

Third-instar larvae crawled up the side of the plastic container with the larval rearing medium and pupated directly onto a thin layer of sterilised sand. The sand was sieved daily over 5 days to allow for synchronous adult eclosion in each collected batch of pupae.

To evaluate optimal egg production, each cage was populated with different volumes of pupae at an assumed 1:1 male to female ratio. Tested densities ranged between 2,000 and 24,000 pupae per cage (at 2000 intervals), and each density was replicated seven times. Eggs were collected daily between days 5 and 24 post adult eclosion, and their numbers were estimated volumetrically, accounting for 22,100 eggs per ml (unpublished data/previous study). Based on attained data in our laboratory study, 1 ml of egg volume corresponded to approximately 22,100 eggs. At the end of the collection period, the total egg production per cage was calculated. The pre-oviposition period (time between eclosion and first egg collection) and the period between first and last egg collection in each cage were also recorded.

To evaluate optimal pupal production, different volumes of eggs (0.25-3.00 ml, at 0.25 intervals) were seeded per 1 kg of rearing medium as described above. Six replicates for each egg volume were tested.

### Quality control parameters

#### Egg production

During each egg production experiment, 100-egg samples were taken at 24 h intervals to measure egg-hatch rates. Each sample was placed on a damp filter paper within a 90-mm Petri dish (to avoid desiccation) and allowed to develop to the larval stage. The number of unhatched or damaged eggs was recorded.

#### Pupal production

Daily pupal production was measured volumetrically. The volume (ml), weight (g) and developmental time from egg seeding to the appearance of the first pupae, of each pupal cohort was measured. The total number of pupae obtained from each treatment was determined by dividing the total weight of pupae in each collection by the average weight of one pupa (mean of 100-pupae sample from each pupal collection) (FAO/IAEA/USDA 2014).

#### Statistical Analysis

Comparison of egg production between cages with different pupal densities and pupal production between different egg densities were performed using One-way ANOVA with post-hoc Tukey HSD test. P-values of ≤ 0.05 were considered statistically significant. All statistical analyses were performed using the Minitab 16 software.

## Results

### Egg production

Mating-based insect control programmes require mass-rearing with high fecundity rates and the production of millions of insects per week. Female fecundity is influenced by the cage’s size (resting surface area), oviposition surface areas, air circulation inside the cage, rearing medium type, and environmental factors such as temperature and humidity. To evaluate optimal egg production, we populated medfly adult cages with different pupal densities, while retaining all other parameters constant.

Figure 1 shows the total egg production for each pupal density tested. Significant differences in total egg production were observed across the tested density range (F=264.65; d.f.= 11; P< 0.001) (Figure 1). For densities between 2,000 and 18,000 pupae per cage, a correlation between pupal density and total egg production was observed, while higher pupal densities (>18,000 pupae per cage) negatively affected total egg production. The coefficient of determination (R²) between pupal volume and total egg production was 96% (y=−2.36×^2^ + 34.82x + 2.94; R^2^=0.9602) indicating that the curve provides a good fit to the data. These data show that peak egg production per cage can be reached with pupal densities ranging between 14,000 and 18,000 pupae per cage, with pupal densities outside of this range yielding significantly lower numbers of eggs.

**Figure 1:**
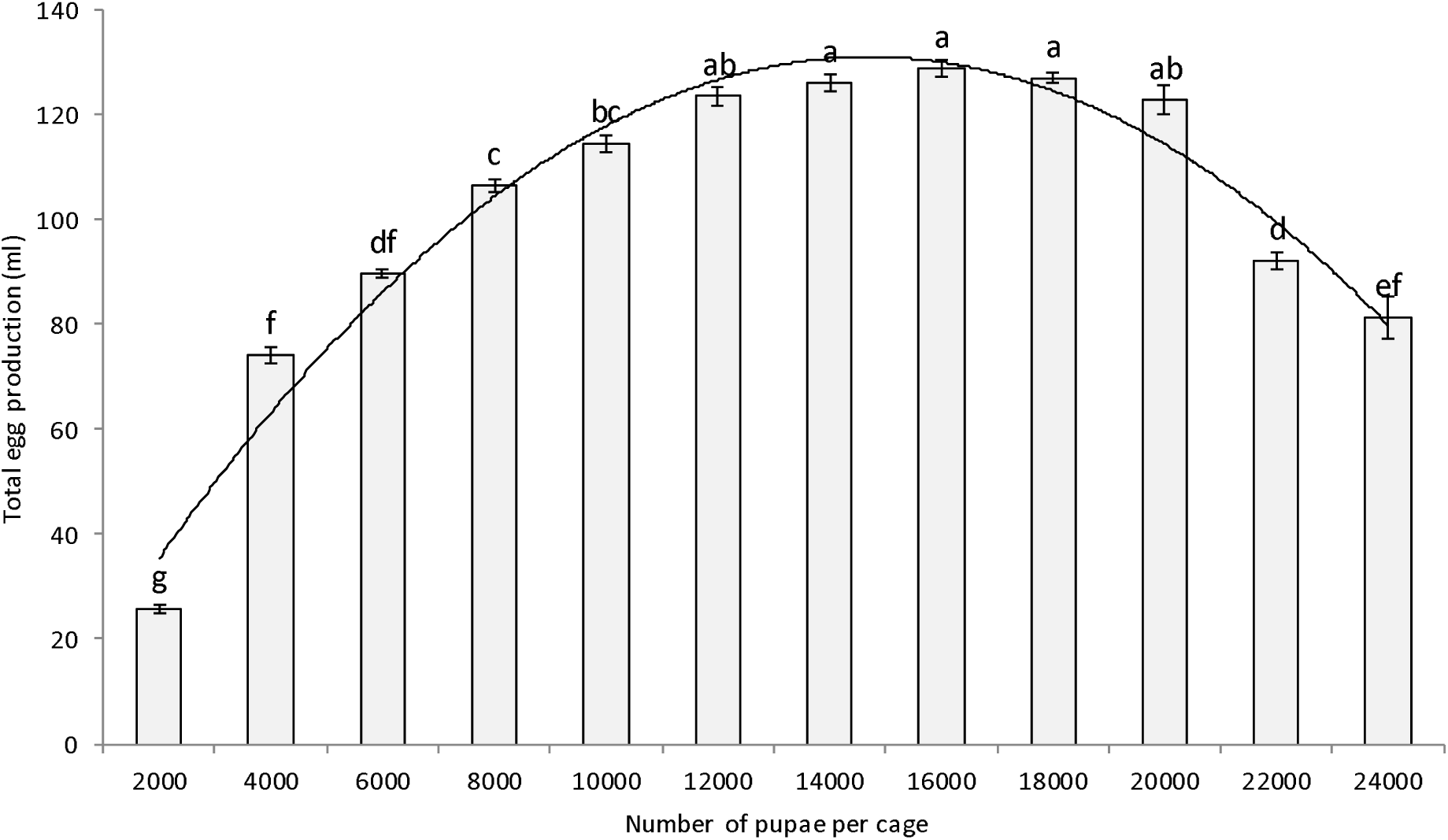
Comparative analysis on the total volume of eggs (ml) obtained from cages populated with varying amounts of OX3864A pupae (mean ± SD; n = 7). Bars superscripted with different letters are significantly different according to ANOVA and Tukey’s HSD test (P>0.05).

Female density per cage affects total egg production per cage, however this value may be inversely proportional to the highest number of eggs laid by female flies, when they subsist under more relaxed cage conditions. To test this hypothesis, we calculated the egg number per female per day, by dividing the number of eggs produced daily by half the number of pupae (assuming a 1:1 male : female eclosion ratio) placed in the mass-rearing cages.

Regarding the daily oviposition rate per female per cage, statistically significant differences were observed between pupal densities (F = 580.32; d.f. = 11; P < 0.001). With the exception of one density (4,000 pupae; Figure 2), a negative correlation was identified between the number of females per cage and the number of eggs produced per female. The coefficient of determination (R²) for total oviposition rate per female between pupal densities was 89%.

**Figure 2:**
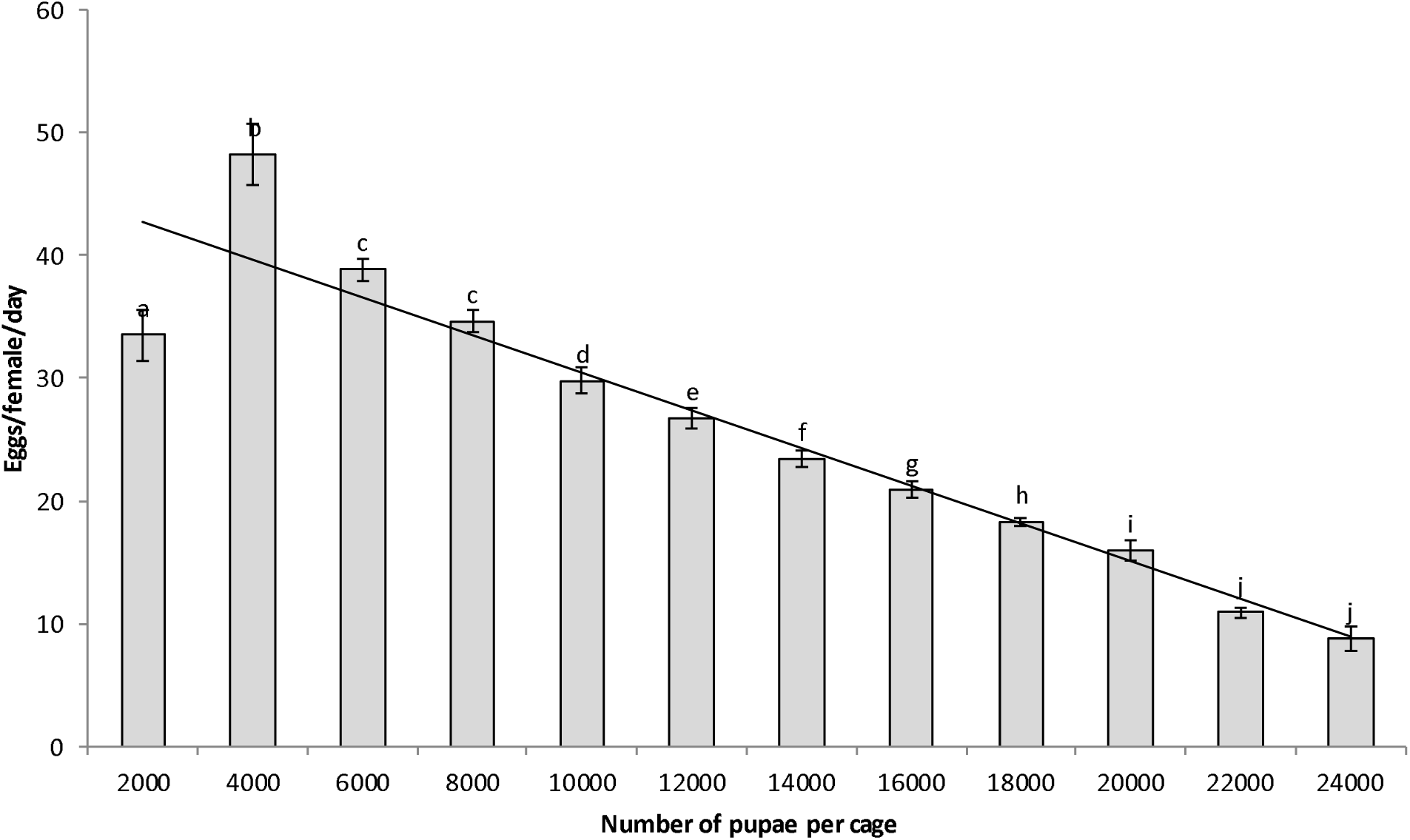
Estimated number of eggs produced per female per day, within each cage populated with different pupal densities (mean ± SD; n = 7). Bars superscripted with different letters are significantly different according to ANOVA and Tukey’s HSD test (P > 0.05).

No statistically significant difference between pupal densities was observed for other egg production parameters such as a) the time between first adult emergence and sexual maturation (as determined by the start of oviposition) (5.01±0.28 days; F = 0.76; d.f. = 11; P = 0.674), b) the period during which oviposition was observed (16±0.43 days; F = 0.76; d.f. = 11; P = 0.677), and c) the mean egg-hatch rate (88.7±1.37%; F = 1.55; d.f. = 11; P = 0.133). High-quality eggs (determined by hatch rates > 85%) were produced for an average of 2 weeks, until approximately 3 weeks post adult eclosion.

Daily egg volumes (ml), and therefore egg production, for each cage treatment are shown in Figure 3. Generally, for all cage treatments, eggs were not produced beyond 23 days post eclosion, however a decline in egg production was observed in most cage treatments by 20 days post eclosion. Egg production peaked on days 10-13 post eclosion for cages with lower pupal densities (2000-12,000 pupae / cage) (Figure 3), except for one (density: 2,000 pupae, peak production: days 6 and 7 post eclosion). For higher pupal densities (14,000-20,000 pupae per cage), peak egg production was observed on days 7-10 post eclosion (Figure 3). For the highest pupal densities (22,000 and 24,000 pupae per cage), increased egg volumes were obtained at days 10-11 post eclosion (Figure 3).

**Figure 3:**
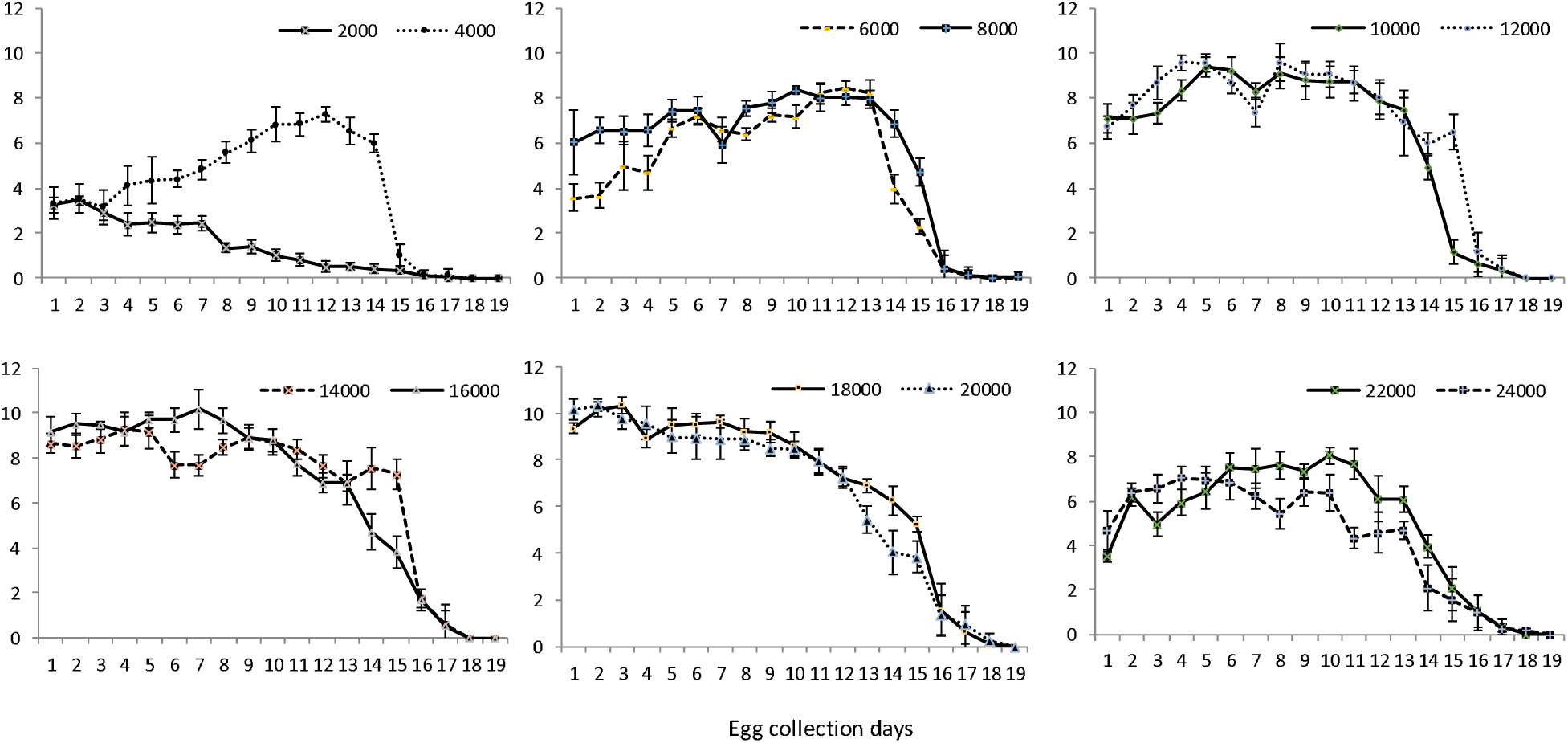
Average daily fecundity of OX3864A females in cages populated with variable pupal densities (mean ± SD; n=7). Egg collection commenced at day 5 post adult eclosion for all the cages.

### Pupal production

The larval development time in the presence of tetracycline (Tet+; F = 74.84; d.f. = 11; P < 0.001), was significantly higher than in the absence of the antibiotic (Tet-; F = 93.00; d.f. = 11; P < 0.001) (Figure 4; Panels A and B, respectively). This was expected as the female-specific self-limiting trait affects females at early stages, largely at the pre-pupal developmental stage. For both Tet+ and Tet-treatments, a direct correlation between pupal production (ml) and egg density was observed up to a density of 1.5 ml eggs/kg. Higher egg densities resulted in fewer pupae produced. The coefficient of determination (R²) between pupal production and egg densities per kg were 92% and 98%, for Tet+ and Tet-, respectively. Our data indicate that egg density of 1.25 ml/kg on Tet+ rearing medium resulted in the highest number of pupae, while egg densities of 1.5-2 ml/kg on Tet-rearing medium yielded the highest quantity of male-only pupae.

**Figure 4:**
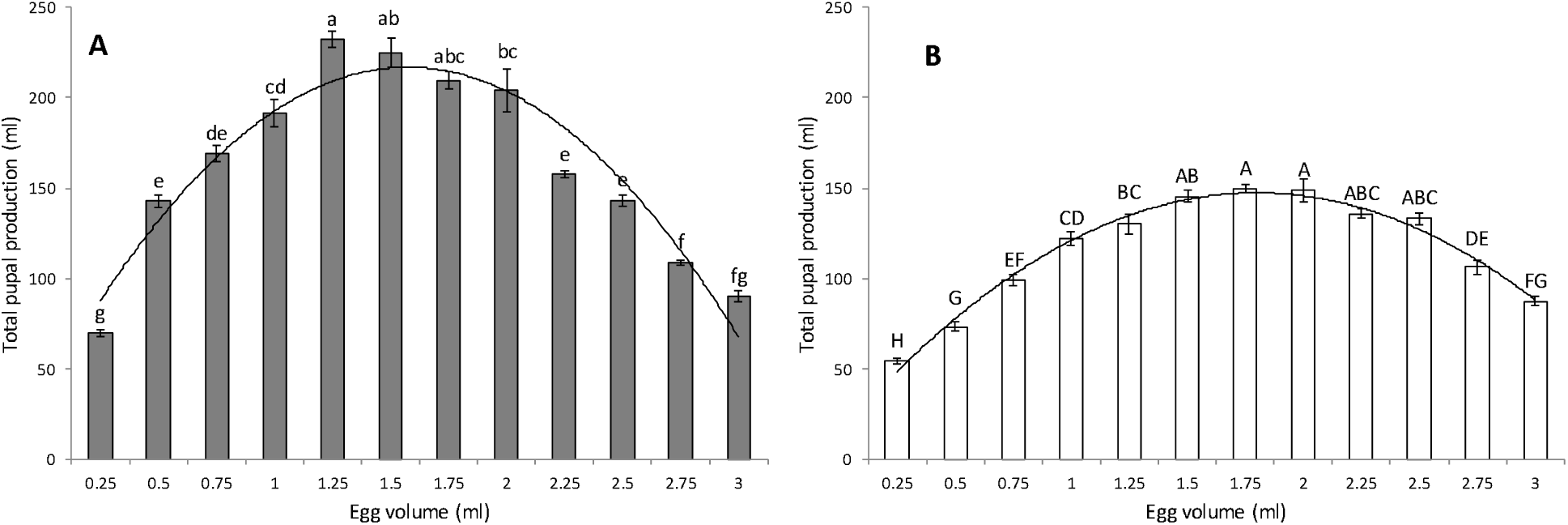
Comparison of pupal production (ml) for different volumes of OX3864A eggs seeded into 1 kg larval rearing medium, under permissive (Tet+) **(A)** and restrictive (Tet-) **(B)** conditions. Mean volume of pupae produced is shown for each initial egg density (mean ± SE; n = 6). Mean values labelled with the same letter are not significantly different according to ANOVA and Tukey’s HSD test (P > 0.05).

Significant differences were observed in total pupal production and pupal weight from different volumes of eggs seeded into 1 kg of (Tet+) rearing medium (F = 39.66; d.f. = 11; P <0.001) (Table 1). The best pupal production values resulted from an egg density of 1.25 ml/kg larval rearing medium. However, the heaviest pupae originated from egg densities between 0.25 ml/kg and 0.75 ml/kg, (average weight=9 mg per pupa). Significant differences in larval development time have also been observed (F = 4.53; d.f = 11; P <0.001), with higher egg densities leading to longer times for pupal recovery. By contrast, egg to pupa recovery (F = 122.05; d.f = 11; P <0.001) and pupal weight per 100 pupae (F = 13.53; d.f = 11; P <0.001) were negatively affected by higher egg densities. However, no significant difference in the duration of pupal collection (4 days for all egg densities) was found, (F = 1; d.f = 11; P = 0.453). Male only pupal production was highest at the density of 2 ml eggs per 1 kg of (Tet-) rearing medium (Table 2). For all egg densities, no significant difference was observed for the time required for complete larval development (10 days; F = 0.9; d.f = 11; P = 0.574), and for the duration of pupal collection (4 days; F = 1.45; d.f = 11; P = 0.174). Higher egg densities negatively affected egg to pupae recovery rates (F = 1.45; d.f = 11; P <0.001) and mean pupal weight per 100 pupae (F = 13.53; d.f = 11; P <0.001). The highest pupal recovery rate of 59.69±1.99 was obtained at a density of 0.25 ml/kg of larval rearing medium. The lowest pupal recovery rates of 10.20±0.56 and 8.00±0.2 were recorded at the egg densities of 2.75 ml/kg and 3.0 ml/kg of Tet-rearing medium, respectively.

**Table 1:**
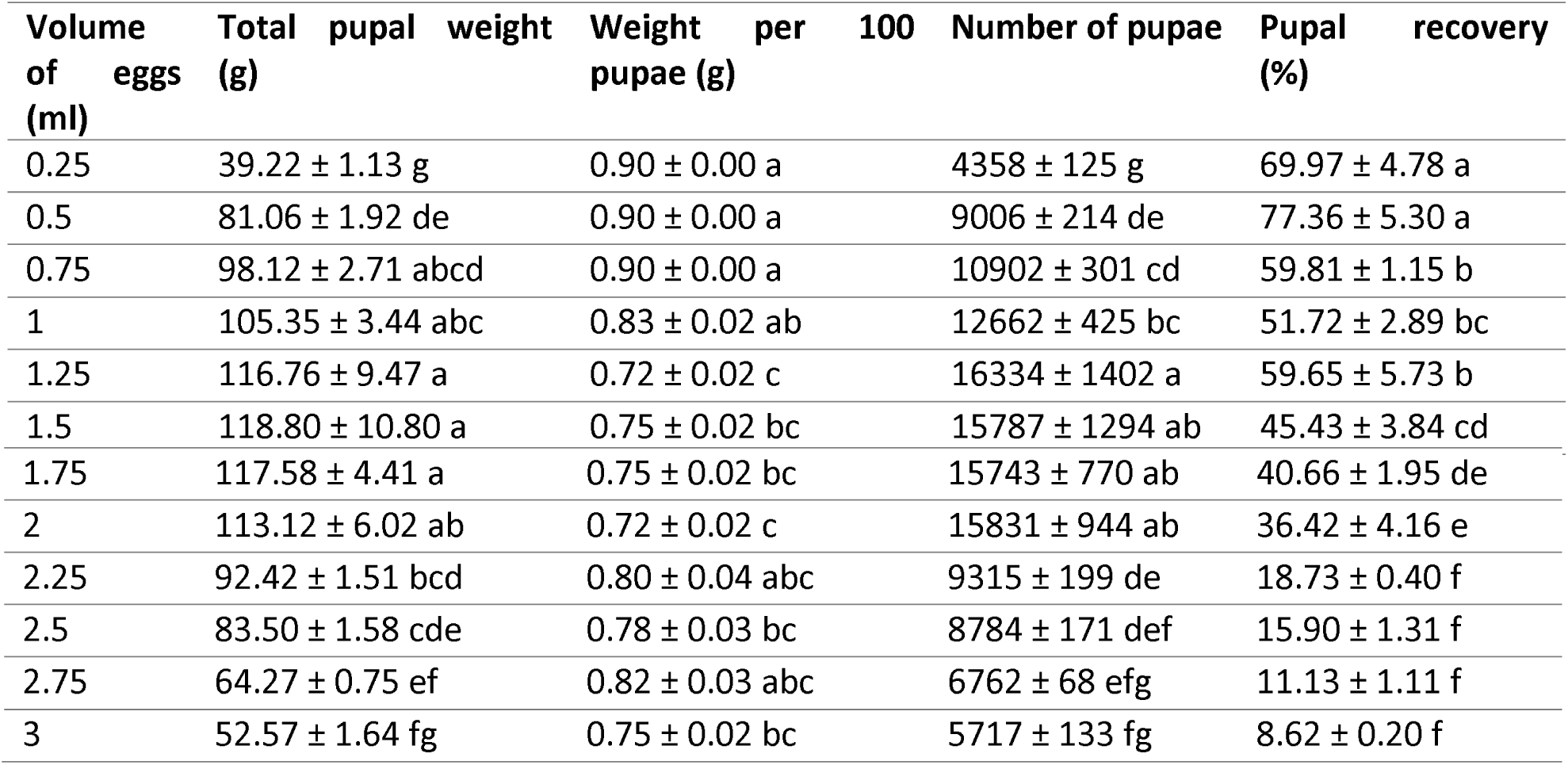
Comparative pupal recovery rates (%) and quality control parameters for different volumes of OX3864A eggs seeded into 1 kg rearing medium, under permissive conditions (Tet+) (mean ± SE; n = 6). Data followed by the same letter, in the same column, do not differ significantly according to ANOVA and Tukey’s HSD test (P>0.05).

**Table 2:**
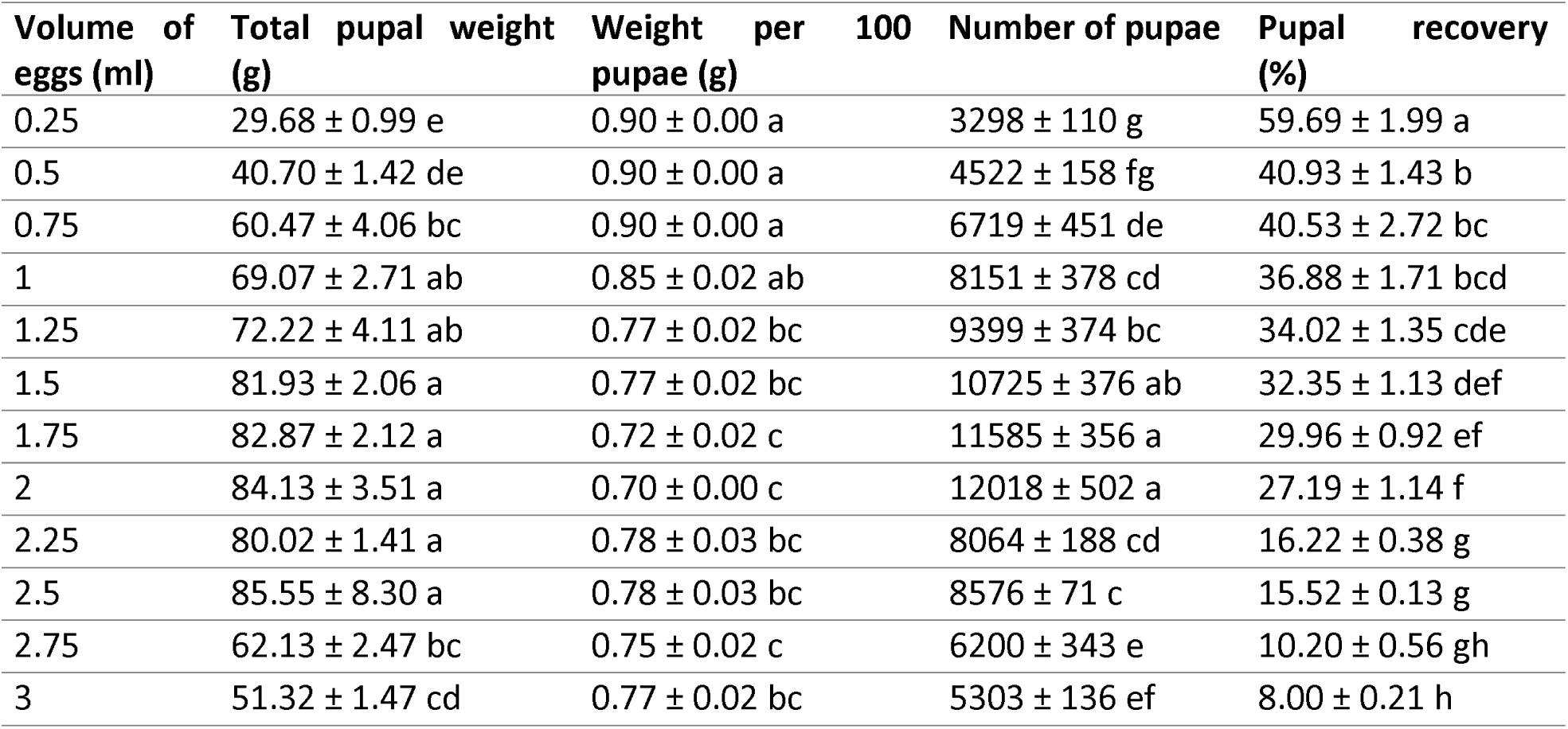
Comparative pupal recovery rates (%) and quality control parameters for different volumes of OX3864A eggs seeded into 1 kg rearing medium, under restrictive conditions (Tet-) (mean ± SE; n=6). Data followed by the same letter, in the same column, do not differ significantly according to ANOVA and Turkey’s HSD test (P>0.05).

For each egg density, a sample of 100 pupae was tested for adult eclosion on (Tet+) and (Tet-) rearing medium (Table 3). For the Tet+ rearing medium, no significant differences were observed for male (F = 0.35; d.f = 11; P = 0.969), female (F = 1.2; d.f = 11; P = 0.305), or total adult (F = 1,44; d.f = 11; P = 0.178) eclosion rates, as well as pupal mortality rates (9.58± 0.508%; F = 1.23; d.f = 11; P = 0.271). For the Tet-rearing medium, male (F= 0.79; d.f = 11; P = 0.648), female (no females for all densities) and total adult (F = 0.79; d.f = 11; P = 0.648) eclosion rates, as well as pupal mortality rates (7.88± 0.797%; F = 0.72; d.f = 11; P = 0.713) were also similar.

**Table 3:** Comparative male and female adult eclosion rates when different volumes of OX3864A eggs were seeded into 1 kg rearing medium, under permissive (Tet+) and restrictive (Tet-) conditions (Mean ± SE; n=6). Data followed by the same letter, in the same column, do not differ significantly according to ANOVA and Turkey’s HSD test (P>0.05).

| Egg<br>Volume<br>(ml) | (Tet+) Larval Rearing medium |  | (Tet-) Larval Rearing medium |  |
| --- | --- | --- | --- | --- |
|  | Male<br>emergence (%) | Female<br>emergence (%) | Male emergence<br>(%) | Female<br>emergence (%) |
| 0.25 | 46.17 ± 1.33 a | 47.17 ± 1.30 a | 92.83 ± 1.05 a | 0.00 ± 0.00 a |
| 0.5 | 46.33 ± 0.56 a | 46.67 ± 0.49 a | 89.50 ± 2.40 a | 0.00 ± 0.00 a |
| 0.75 | 44.33 ± 2.40 a | 47.50 ± 1.28 a | 94.33 ± 1.41 a | 0.00 ± 0.00 a |
| 1 | 43.00 ± 2.89 a | 49.00 ± 2.34 a | 92.00 ± 1.59 a | 0.00 ± 0.00 a |
| 1.25 | 43.33 ± 1.93 a | 46.00 ± 1.57 a | 91.00 ± 1.59 a | 0.00 ± 0.00 a |
| 1.5 | 42.33 ± 3.36 a | 42.00 ± 2.94 a | 90.00 ± 2.67 a | 0.00 ± 0.00 a |
| 1.75 | 43.00 ± 2.07 a | 45.17 ± 1.64 a | 92.33 ± 1.36 a | 0.00 ± 0.00 a |
| 2 | 42.67 ± 2.78 a | 42.83 ± 2.85 a | 94.67 ± 1.05 a | 0.00 ± 0.00 a |
| 2.25 | 43.00 ± 3.18 a | 50.00 ± 3.53 a | 91.17 ± 2.77 a | 0.00 ± 0.00 a |
| 2.5 | 43.17 ± 3.09 a | 49.33 ± 2.97 a | 89.67 ± 2.56 a | 0.00 ± 0.00 a |
| 2.75 | 46.00 ± 2.65 a | 47.67 ± 1.69 a | 93.00 ± 1.46 a | 0.00 ± 0.00 a |
| 3 | 42.17 ± 3.19 a | 46.17 ± 1.96 a | 91.83 ± 1.76 a | 0.00 ± 0.00 a |

For all egg densities on Tet+ and Tet-rearing medium, the first day of pupal collection yielded the highest number of pupal volume, while the pupae yield from fourth day was very low.

## Discussion

Previous SIT programmes worldwide have been successful at locally suppressing or eradicating pest populations of medfly (Enkerlin et al. 2015). However, their cost effectiveness may be compromised by the irradiation employed to sterilise males and the resulting decrease in mating competitiveness. OX3864A does not need irradiating and may present a more cost-effective alternative to pest population suppression, provided mass rearing of the strain shows comparable standards to SIT strains.

Under the conditions tested in this study and at a cage density of 18,000 pupae, mean egg production for OX3864A doubled the reported egg production for the SIT strain VIENNA-8 *tsl* (Temperature Sensitive Lethal, genetic sexing strain; Neto et al., 2012). Furthermore, Rempoulakis et al. (2016) also reported lower egg production for three *tsl* strains (VIENNA-8, VIENNA-8.Sr^2^ and VIENNA 8-1260). Our data also showed egg hatching rates across all experimental treatments were 27-48% higher than the rates reported for VIENNA-8 *tsl* strain by Neto et al. (2012) and Rempoulakis et al. (2016). Caceres et al. (2002) showed that the main cost difference between rearing a *tsl*-based Genetic sexing strain and a wild type strain is directly linked to their respective egg production efficiency. Taken together, the higher egg yields, better egg hatch rates and the high stability of the OX3864A Oxitec medfly strain translate in the reduction of the cost by 59% in producing 100 million males per week when compared to the *tsl* (Table 4), a significant cost savings in a mass-rearing setting. Our calculations were based on Caceres et al. (2002) description of the relative efficiencies of a *tsl* SIT genetic sexing strain including staffing requirements, consumables and equipment as well as the rental of the facility.

**Table 4:**
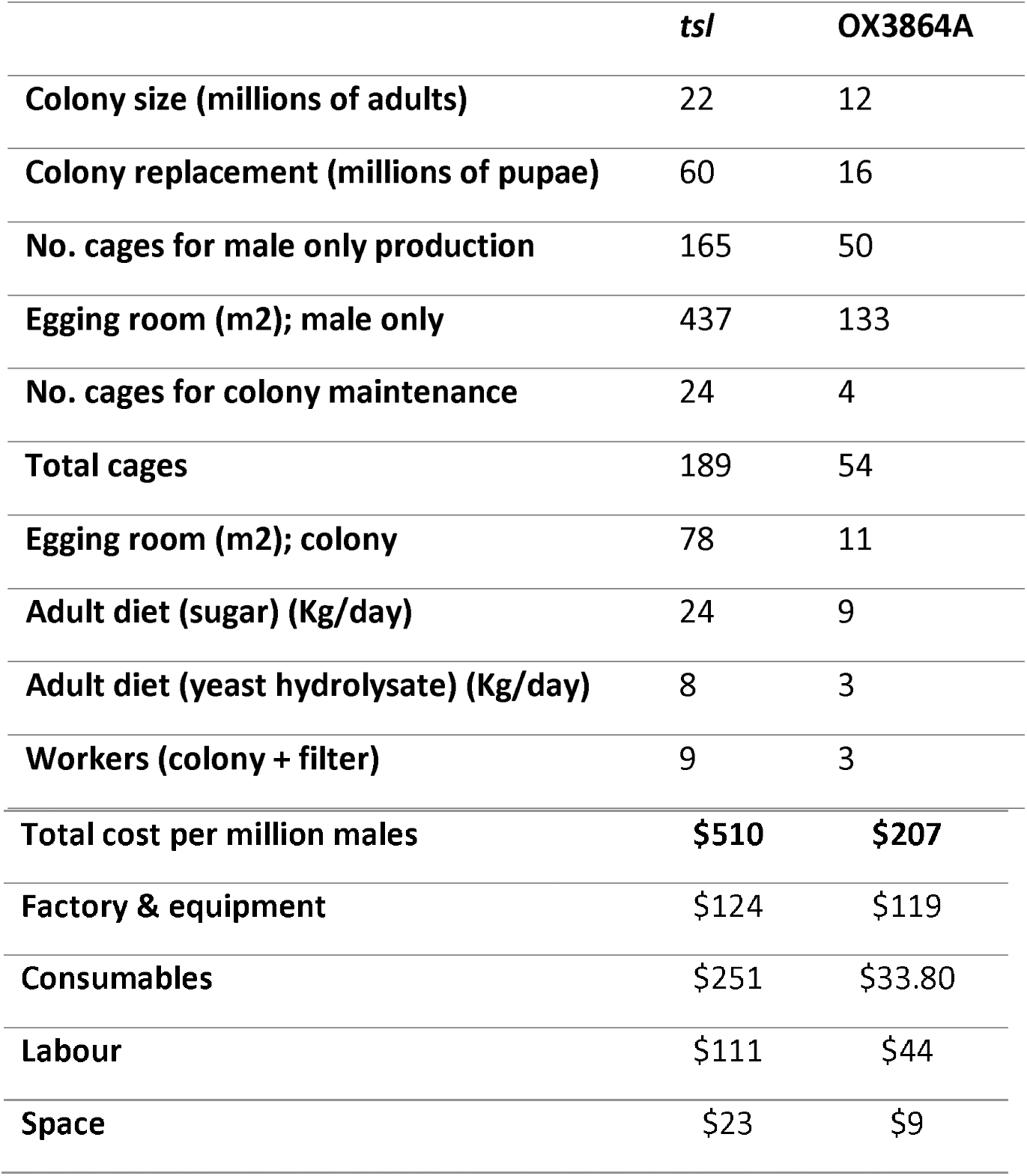
Comparison of rearing requirements and rearing cost between *tsl* and OX3864A to produce 100 million medfly males per week. (Caceres et al. 2002, personal communication with Oxitec Ltd modelling team for OX3864A cost analysis)

For maximum efficiency, egg collections should be timed to ensure a high number of fertilised eggs in the shortest possible time. Taking into account factors such as sexual maturation of female medflies (∼day 3 post eclosion; Chapman et al. 1998, Lance et al. 2000, Lux et al. 2002, Robinson et al. 2002), mating period (∼days 3-5 post eclosion; Gordillo 1996), peak egg production and high egg hatch rates (>85%), a recommended period for egg collections in all cage densities is between days 5-19 post adult eclosion. Egg collections performed after 9 days showed increasing female mortality within the cages and decreasing female fecundity suggesting reduced cost efficiency for a mass-rearing system. This optimal egg collection during two weeks, which is comparable to the VIENNA-8 strain (Hamden et al. 2013), ensures a steady and uninterrupted supply of sterile OX3864A males during a suppression programme.

We have previously shown the capacity of the self-limiting OX3864A strain to suppress wild type population of medfly in cage studies (Leftwich et al. 2014). With the present study, we have investigated mass-rearing parameters related to strain propagation and production of a male-only cohort for field release, in a mating-based operational programme. Our data demonstrate the potential of OX3864A strain to be mass reared successfully and potentially more cost-effectively than previous SIT strains. Further in-country analysis will be required if the self-limiting strain OX3864A becomes part of an operational programme for medfly control.

## Acknowledgements

We thank Zoubida Id-Zzaouit for technical assistance and guidance during experiments, Salah Salmati and Said Akhzam for rearing medium preparation, cage maintenance and egg/pupal collections and measurements. We also thank Dr. Simon Warner and Dr. Enca Martin-Rendon for their constructive comments on this manuscript.

